# Transcriptomic responses to warming and cooling of an Arctic tundra soil microbiome

**DOI:** 10.1101/599233

**Authors:** Morten Dencker Schostag, Muhammad Zohaib Anwar, Carsten Suhr Jacobsen, Catherine Larose, Timothy M. Vogel, Lorrie Maccario, Samuel Jacquiod, Samuel Faucherre, Anders Priemé

**Author notes:** Corresponding author: Anders Priemé.

## Abstract

**Background:** Arctic surface soils experience pronounced seasonal changes in temperature and chemistry. However, it is unclear how these changes affect microbial degradation of organic matter, nitrogen cycling and microbial stress responses. We combined measurements of microbiome transcriptional activity, CO_2_ production, and pools of carbon and nitrogen to investigate the microbial response to warming in the laboratory, from −10 °C to 2 °C, and subsequent cooling, from 2 °C to −10 °C, of a high Arctic tundra soil from Svalbard, Norway.

**Results:** Gene expression was unaffected by warming from −10 °C to −2 °C and by cooling from −2 °C to −10 °C, while upon freezing (2 °C to −2 °C) a defense response against oxidative stress was observed. Following modest transcriptional changes one day after soil thaw, a more pronounced response was observed after 17 days, involving numerous functions dominated by an upregulation of genes involved in transcription, translation and chaperone activity. Transcripts related to carbohydrate metabolism and degradation of complex polymers (e.g. cellulose, hemicellulose and chitin) were also enhanced following 17 days of soil thaw, which was accompanied by a four-fold increase in CO_2_ production. In addition, anaerobic ammonium oxidation and turnover of organic nitrogen were upregulated. In contrast, nitrification, denitrification and assimilatory nitrate reduction were downregulated leading to an increase in the concentration of soil inorganic nitrogen.

**Conclusion:** the microorganisms showed negligible response to changes in sub-zero temperatures and a delayed response to thaw, which after 17 days led to upregulation of soil organic matter degradation and enhanced CO_2_ production, as well as downregulation of key pathways in nitrogen cycling and a concomitant accumulation of inorganic nitrogen available for plants.

## Background

Permafrost soil systems cover ~20 % of the non-glaciated land surface in the Northern Hemisphere (Zhang et al., 1999). These northern soils are currently affected by larger temperature increases than lower latitude soils (Duarte et al., 2012) and this is predicted to continue during future climate change (Pachauri et al., 2014), especially during winter (Collins et al., 2013). Due to enhanced warming, permafrost soil is thawing and the seasonally thawed active layer soil on the surface has deepened over the past decades (Elberling et al., 2013) and this will likely worsen in the future (Anisimov et al., 1997; Lawrence & Slater, 2005). Permafrost soils contain ca. 1700 Pg organic carbon (Tarnocai, 2009) with the majority found in the upper soil profile (Hugelius et al., 2014). Parts of this frozen carbon pool might be transformed to greenhouse gasses by a high diversity of microorganisms (Jansson & Taş, 2014; Mackelprang et al., 2016) when permafrost soils thaw and become active layer soils (Schuur et al., 2015). In addition, nitrogen cycling in active layer soil may be affected by warming (Blok et al., 2018; Phillips et al., 2019). Most Arctic terrestrial ecosystems are characterized by nitrogen limitation (Tamm, 1991; Elser et al., 2007) and generally receive low amounts of atmospheric nitrogen deposition (<2 kg N ha^−1^ y^−1^) (Dentener et al., 2006). Not only transformation of soil organic matter (Chen et al., 2014), but also plant growth is closely linked to soil nutrient availability, and the current ‘greening’ of the Arctic (Elmendorf et al., 2012) may over time be slowed down by limited plant-available soil nitrogen. Thus, the transition from permafrost to active layer soil and the projected increase in active layer soil temperature in coming decades may be involved in several feedbacks on global warming (Pachauri et al., 2014), notably through changes in carbon and nitrogen cycling processes.

Active layer soils are highly dynamic environments with large annual amplitudes in temperature as well as water and nutrient availability. Future increases in temperature and reductions in snowfall will not only affect soil temperature and its variability/stability, but also the number of freeze-thaw cycles for surface soils (Williams et al., 2015), which influence microbial activity and survival (Buckeridge et al., 2013), and hence, greenhouse gas emission rates (Priemé et al., 2001). Despite sub-zero temperatures, low water and nutrient availability, and freeze-thaw cycles during winter, a high diversity of microorganisms can be found in active layer soils (Chu et al., 2010; Tveit et al., 2013; Schostag et al., 2015).

Microbial activity in permafrost soil was observed down to −39 °C (Panikov et al., 2006) and degradation of cellulose at −4 °C (Segura et al., 2017), while growth of bacteria isolated from active layer soil was recorded at −15 °C (Mykytczuk et al., 2013) and DNA replication at −20°C (Tuorto et al., 2014). Microbial activity at these low temperatures involves a wide range of molecular mechanisms including stress responses (D’Amico et al., 2006; Bakermans et al., 2012). Stress protection is costly to microorganisms and a trade-off exists between stress response and non-stress related activities (Hõrak & Tamman, 2017). Thus, microorganisms are known to downregulate many non-stress genes as part of their stress response (Horn et al., 2007) and this may influence, e.g., the initiation of soil organic matter transformation and cycling of inorganic nitrogen following thaw of active layer soils.

Although a few metagenomic studies have highlighted potential microbial metabolic pathways and survival mechanisms at low temperatures in permafrost and active layer soil (Yergeau et al., 2010; Mackelprang et al., 2011; Lipson et al., 2013), we still lack information on actual activity patterns showing which genes are expressed under these conditions. Furthermore, interpretation of DNA-based data is hampered by the persistence of extracellular DNA in cold soils (Willerslev et al., 2004). Indeed, up to 40 % of DNA isolated from surface soil is extracellular or from cells that are no longer intact, hence their DNA-based signal in cold soils might not be correlated to actual microbial activity (Carini et al., 2016). To our knowledge, only two studies have investigated gene expression in soil from cold environments at sub-zero temperatures using mRNA-based techniques (Coolen & Orsi, 2015; Hultman et al., 2015). Hultman et al. (2015) used a multi-omics approach and reported that cold shock protein genes were transcribed at a higher abundance in the colder permafrost compared to the warmer active layer. This study only involved a single time point under frozen conditions, whereas (Coolen & Orsi, 2015) compared gene transcription at a single time point before and one after thawing permafrost soil, revealing an enhanced transcription of genes related to DNA repair functions and enhanced biofilm formation in frozen soil as compared to after thawing.

At present, the soil microbial responses to i) temperature change at sub-zero temperature, ii) thawing, and iii) freezing are not fully understood. Thus, we conducted a laboratory study simulating a short Arctic spring, summer and autumn, where we investigated the microbial expression of mRNA, CO_2_ production, and soil carbon and nitrogen pools at different time points during warming of a frozen active layer soil followed by thawing, re-freezing and cooling. We hypothesized that microbial transcription of genes involved in i) degradation of soil complex organic matter (e.g. lignocellulose and chitin), ii) mineralization of soil organic nitrogen, and iii) cycling of inorganic soil nitrogen, will all increase during warming, especially upon soil thaw, and subsequently decrease during re-freezing. We also hypothesized that transcription of genes involved in microbial stress responses increases upon iv) thawing and v) freezing, and that vi) thaw-induced microbial stress responses postpone transcription of non-stress related genes.

## Results

### Soil parameters, CO_2_ production and RNA concentration

The incubation temperature and sampling points are depicted in Fig. 1. Soil characteristics measured prior to incubation is in Supporting Information Table S1, while data on soil chemical parameters measured during the experiment are in Table 1. pH was within 7.4 and 7.6 during the experiment. DOC and DON declined markedly (*P* < 0.05) when soil temperature was above zero, while nitrate concentration increased between W_2°C_ and C_2°C_ (*P* < 0.05) before receding to the initial level. Ammonium concentration increased between C_2°C_ and C_−6°C_ (*P* < 0.05). Carbon dioxide production rates increased significantly (*P* < 0.05) from 66 ± 15 ng CO_2_ g^−1^ dwt soil h^−1^ (average ± standard error of the mean, n = 4) at W_−6°C_ to 105 ± 16 ng CO_2_ g^−1^ dwt soil h^−1^ at W_2°C_ and 423 ± 5 ng CO_2_ g^−1^ dwt soil h^−1^ at C_2°C_. With a methane uptake of 9.6 pg CH_4_ g^−1^ dry soil h^−1^ at W_2°C_, the soil changed from a net methane sink to a net source of methane at W_2°C_ with emission of 13.1 pg CH_4_ g^−1^ dry soil h^−1^. Only negligible rates of nitrous oxide emissions were observed. Total RNA concentration after DNase treatment was similar among all samples (ANOVA with a post-hoc Tukey’s HSD correction test; *P* > 0.05; see Schostag et al., 2019).

**Table 1.** Soil physiochemical parameters at different incubation temperatures (W: warming; C: cooling). Data are average ± standard error of the mean, n = 5. Different superscript letters indicate that samples are significantly different (pairwise comparisons between values within each column, ANOVA with Tukey’ HSD posthoc test). DOC: dissolved organic carbon; DON: dissolved organic nitrogen.

|  | DOC | DON | NH<sub>4</sub><sup>+</sup> | NO<sub>3</sub><sup>-</sup> | pH |
| --- | --- | --- | --- | --- | --- |
| | ($\mu\text{g g}^{-1}$ dry soil) | ($\mu\text{g g}^{-1}$ dry soil) | ($\text{ng g}^{-1}$ dry soil) | ($\text{ng g}^{-1}$ dry soil) | |
| W<sub>-6°C</sub> | 174±12<sup>b</sup> | 8.3±0.4<sup>c</sup> | 80±6.5<sup>a</sup> | 82±34<sup>a</sup> | 7.5±0.02<sup>b</sup> |
| W<sub>2°C</sub> | 156±4.6<sup>b</sup> | 8.9±0.3<sup>c</sup> | 90±9.6<sup>a</sup> | 88±14<sup>a</sup> | 7.6±0.04<sup>c</sup> |
| C<sub>2°C</sub> | 58±17<sup>a</sup> | 5.0±0.4<sup>a</sup> | 171±17<sup>a</sup> | 928±199<sup>b</sup> | 7.4±0.03<sup>a</sup> |
| C<sub>-6°C</sub> | 69±7.4<sup>a</sup> | 6.5±0.3<sup>b</sup> | 552±7.4<sup>b</sup> | 153±35<sup>a</sup> | 7.4±0.00<sup>ab</sup> |

**Fig. 1.**
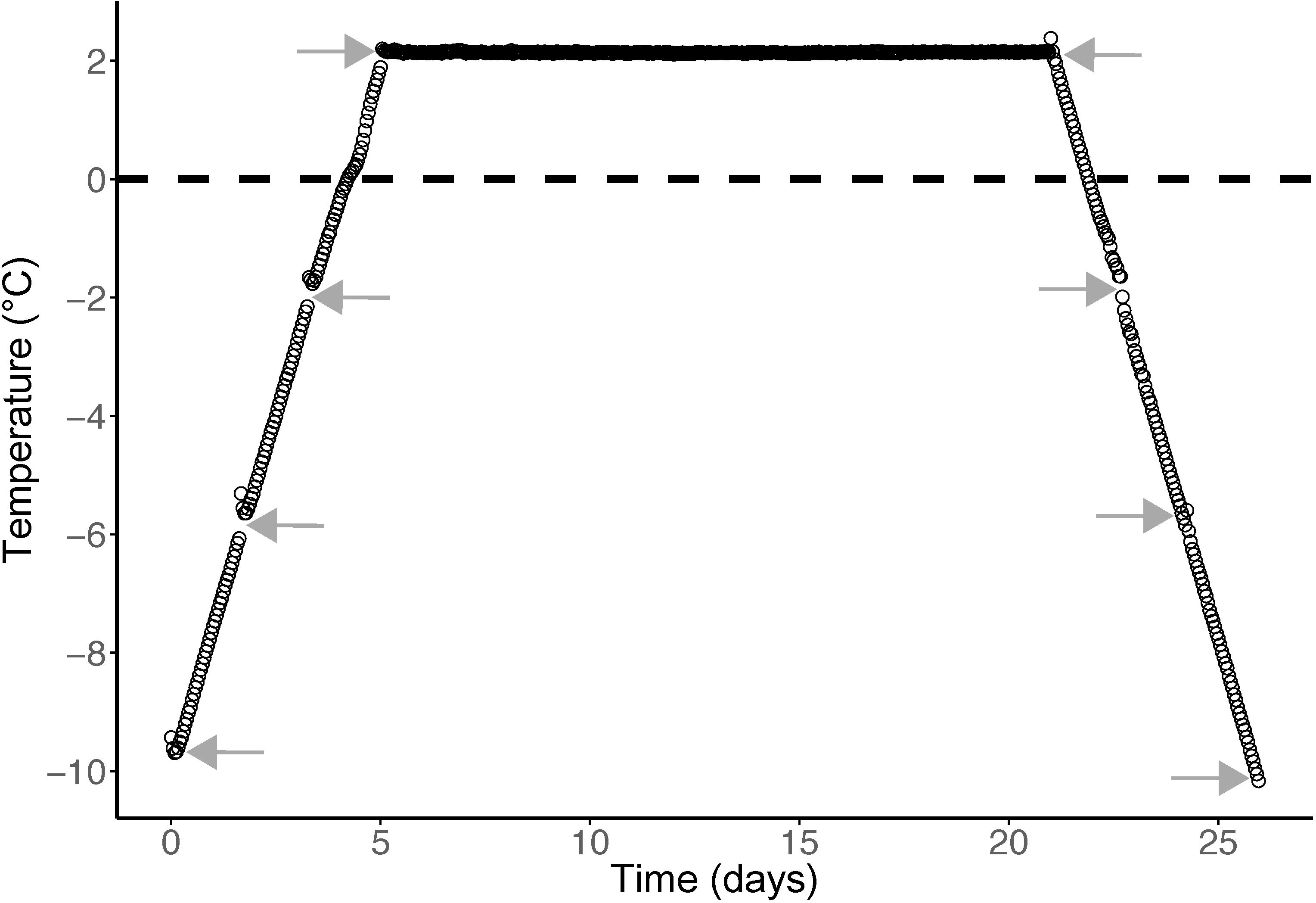
Measured incubation temperature of active layer permafrost samples shown as average of five temperature loggers (black circles); standard error of mean ≤0.1 °C. Arrows indicate time of sampling for RNA isolation.

### Sequencing results

An average (± standard error of the mean) of 35 ± 1.5 million reads per sample (forward and reverse) was obtained from the sequencing. After quality trimming and rRNA removal, the number of reads per sample was reduced to an average of 1.65 ± 0.08 million, representing 4.7 ± 0.12 % of the initial reads. The annotation resulted in an average match of 15,667 ± 953 reads per sample when aligning to the eggNOG database, 53,299 ± 3,334 reads per sample to CAZy database, and 13,616 ± 793 reads per sample to NCycDB, which represented 2.6 %, 8.7 % and 2.2 % of the potential mRNA reads, respectively. The stats for each step from sorting, quality filtering to annotation across each sample are provided in Supporting Information Table S2.

### Overall responses to warming and cooling

The constrained analysis (BGA) grouping replicates under Monte-Carlo simulation revealed a significant, non-random distribution of the groups for the eggNOG, CAZy and NCycDB data (*P* < 10^−6^ for all three data sets), see Fig. 2. The first component of the BGA explained 74.2 % of the variance in the eggNOG data and clearly segregated the warming from the cooling samples, while the second component explained 9.1 % of the variance and separated C_2°C_ from the other cooling samples (Fig. 2A). A similar separation was found for the CAZy and NCycDB data where the first component explained 81.6 % and 80.4 %, respectively, of the variance and clearly segregated the warming from the cooling samples, while the second component explained 8.0 % and 8.8 %, respectively, of the variance and revealed separation of C_2°C_ from the other cooling samples (figs. 2B and 2C). This observation was confirmed by PERMANOVA on Euclidian distance, where 53.9 %, 68.3 % and 58.8 % of the variance was attributed to the treatment (warming samples versus cooling samples, *P* < 10^−6^) in eggNOG, CAZy and NCycDB profiles, respectively. The changing temperature (i.e. warming from −10 °C to 2 °C and cooling from 2 °C to −10 °C) did not show a significant effect, representing 6.3 %, 4.1 % and 5.1 % of the variance in eggNOG, CAZy and NCycDB profiles, respectively.

**Fig. 2.**
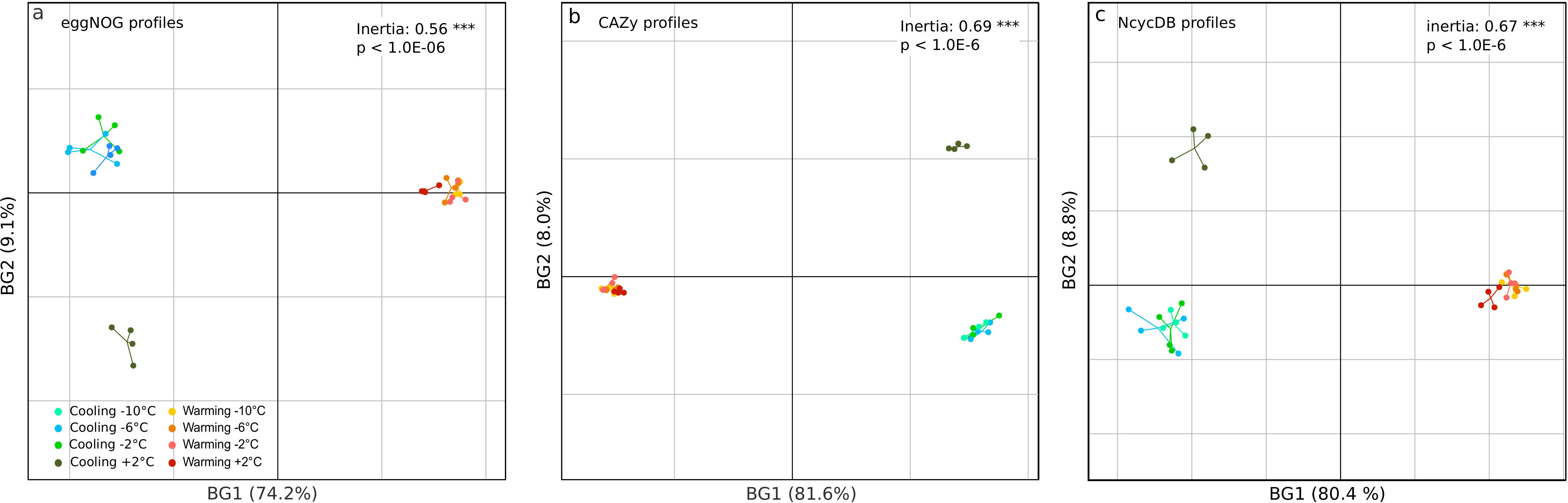
Between-group analysis (BGA) of all functions annotated using (a) eggNOG, (b) CAZy, and (c) NCycDB databases. The figures show constrained principal component analysis (PCA) of the eggNOG, CAZy and NCycDB profiles after applying sample grouping according to replicates. Non-random distribution of the BGA grouping was tested using a Monte–Carlo simulation with 10,000 permutations (*P* < 10^−6^ for all three databases).

### General transcript annotation - alignment against M5nr database and eggNOG annotation

In the total dataset, we detected 652 annotated functions when aligning against the M5nr database and annotating using eggNOG databases. Analysis of differential gene expression was performed using the DESeq2 pipeline with pairwise comparisons between all the samples. The pairwise comparison between the sub-zero samples during the warming or cooling phases revealed only a single eggNOG function that was differentially up or ddownregulated (Table 2). Thus, we decided to combine the two sets of sub-zero samples and perform only three pairwise comparisons, which involved i) all sub-zero warming samples vs. W_2°C_, ii) W_2°C_ vs. C_2°C_, and iii) C_2°C_ vs. all sub-zero cooling samples. A complete list of all gene functions that were significantly up or downregulated at steps i–iii can be found in Supporting Information Data Sheets S1–S3. When comparing all sub-zero warming samples with W_2°C_, 20 out of 652 annotated eggNOG functions encompassing 1.4 % of the total read count (TRC) of mRNA reads were significantly upregulated, while two (representing 0.77 % of TRC) were downregulated.

**Table 2.** Number of eggNOG, CAZy and NCycDB gene categories significantly down or upregulated between the consecutive time points employed in the experiment. W: warming; C: cooling.

|  |  | Downregulated |  |  | Upregulated |  |  |
| --- | --- | --- | --- | --- | --- | --- | --- |
|  |  | eggNOG | CAZy | NCycDB | eggNOG | CAZy | NCycDB |
| Warming | W<sub>-10°C</sub> to W<sub>-6°C</sub> | 1 | 2 | 0 | 0 | 0 | 0 |
|  | W<sub>-6°C</sub> to W<sub>-2°C</sub> | 0 | 0 | 0 | 0 | 0 | 0 |
|  | W<sub>-2°C</sub> to W<sub>2°C</sub> | 5 | 0 | 1 | 8 | 1 | 3 |
| Stable | W<sub>2°C</sub> to C<sub>2°C</sub> | 41 | 62 | 23 | 154 | 82 | 76 |
| Cooling | C<sub>2°C</sub> to C<sub>-2°C</sub> | 8 | 16 | 3 | 31 | 27 | 17 |
|  | C<sub>-2°C</sub> to C<sub>-6°C</sub> | 0 | 0 | 0 | 0 | 0 | 0 |
|  | C<sub>-6°C</sub> to C<sub>-10°C</sub> | 0 | 0 | 0 | 0 | 0 | 0 |

The largest change in expression pattern occurred during the 16 days at 2 °C. Between these two sampling points, a total of 154 (5.7 % of TRC) and 41 (5.6 % of TRC) annotated functions were significantly up and downregulated, respectively. When comparing C_2°C_ with all other cooling samples 52 (2.3 % of TRC) and 11 (2.5 % of TRC) annotated functions were significantly up or downregulated, respectively.

A selection of gene groups with significantly up or downregulated functions annotated in eggNOG is presented in Fig. 3. After 16 days at 2 °C, we observed a pronounced increase in transcripts involved in production of enzymes compared to the frozen state, i.e. transcripts related to transcription (level 2 category K), translation (level 2 category J), and molecular chaperones (level 3 categories COG0071, COG0234, COG0459, and COG0542) (Fig. 3C, D, H). This pattern was also observed for cold shock proteins (level 3 category COG1272) (Fig. 3G), genes involved in prokaryotic motility (level 2 category N) (Fig. 3E), as well as the level 2 categories B (Chromatin structure and dynamics), D (Cell cycle control, cell division, chromosome partitioning), F (Nucleotide transport and metabolism), and U (Intracellular trafficking, secretion, and vesicular transport), while categories L (Replication, recombination and repair) and V (Defense mechanisms) decreased (data not shown).

**Fig. 3.**
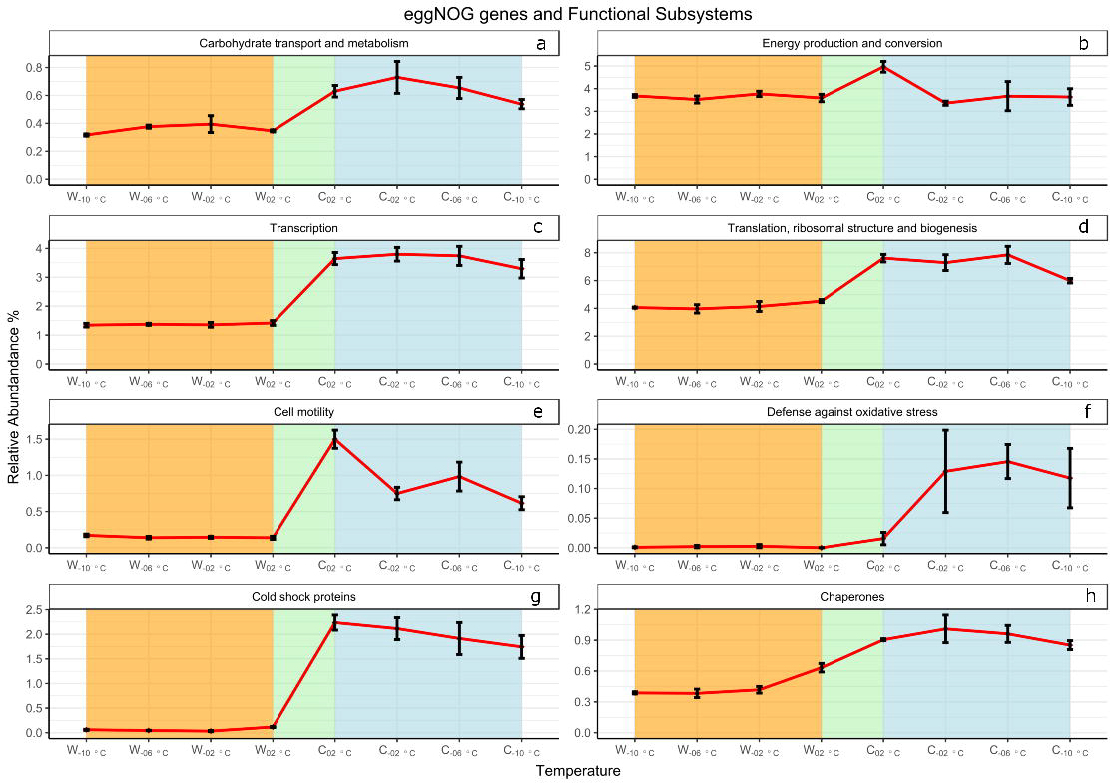
The relative abundance of transcripts related to selected and significantly responding (*P* < 0.05) eggNOG functional groups and genes involved in, a) carbohydrate transport and metabolism, b) energy production and conversion, c) transcription, d) translation, ribosomal structure and biogenesis, e) cell motility, f) defense against oxidative stress, g) cold shock proteins, and h) chaperones. Error bars indicate standard error of the mean.

Following sample freezing, the expression of level 2 categories N (Cell motility) (Fig. 3E), B (Chromatin structure and dynamics) and C (Energy production and conversion) decreased, while expression of genes involved in defense against oxidative stress (level 3 categories COG0753 [catalase] and COG0783 [DNA-binding ferritin-like protein (oxidative damage protectant)]) increased markedly (Fig. 3F).

It should be noted that none of the transcripts annotated in the eggNOG database were related to key genes involved in lignocellulose degradation, inorganic nitrogen cycling or methane cycling *i.e*. genes encoding cellulases, hemicellulases and enzymes with lignolytic activity, and *nifH* (nitrogen fixation), *amoA* (nitrification), *norB*, *norC*, *nirK*, *nirS*, *nos*Z (denitrification), *mrcA* (methane production), and *pmo* (methane oxidation).

### Annotation of transcripts related to degradation of lignocellulose - alignment against CAZy database

When aligning against the CAZy database, we detected a total of 1648 annotated functions (a complete list of all gene functions can be found in Supporting Information Data Sheet S4). Transcripts assigned to CAZy Auxilliary Activity Family 1 or 2 (Levasseur et al., 2013) were related to enzymes with lignolytic activity, e.g. laccases, manganese peroxidases and lignin peroxidases. The abundance of these transcripts decreased following soil thawing and freezing (Fig. 4B). Transcripts assigned to Auxilliary Activity Family 9 or 10 were encoding lytic polysaccharide monooxygenases involved in lignocellulose and chitin degradation (Vaaje-Kolstad et al., 2010; Johansen, 2016) [no reads were assigned to Auxilliary Activity Family 11 and 13], while transcripts assigned to CAZy glycoside hydrolase (GH) families GH5, GH6, GH9, GH44, GH45 and GH48 were presumed to encode cellulases, GH8, GH10, GH11, GH12, GH26, GH28 and GH53 were presumed to encode hemicellulases, and GH18 and GH19 were presumed to encode chitinases. In contrast to the transcripts related to lignolytic activity, the abundance of these four functional groups increased significantly during the 16 days at 2 °C (Figs. 4A, C, D, and Supplementary Information Fig, S1).

**Fig. 4.**
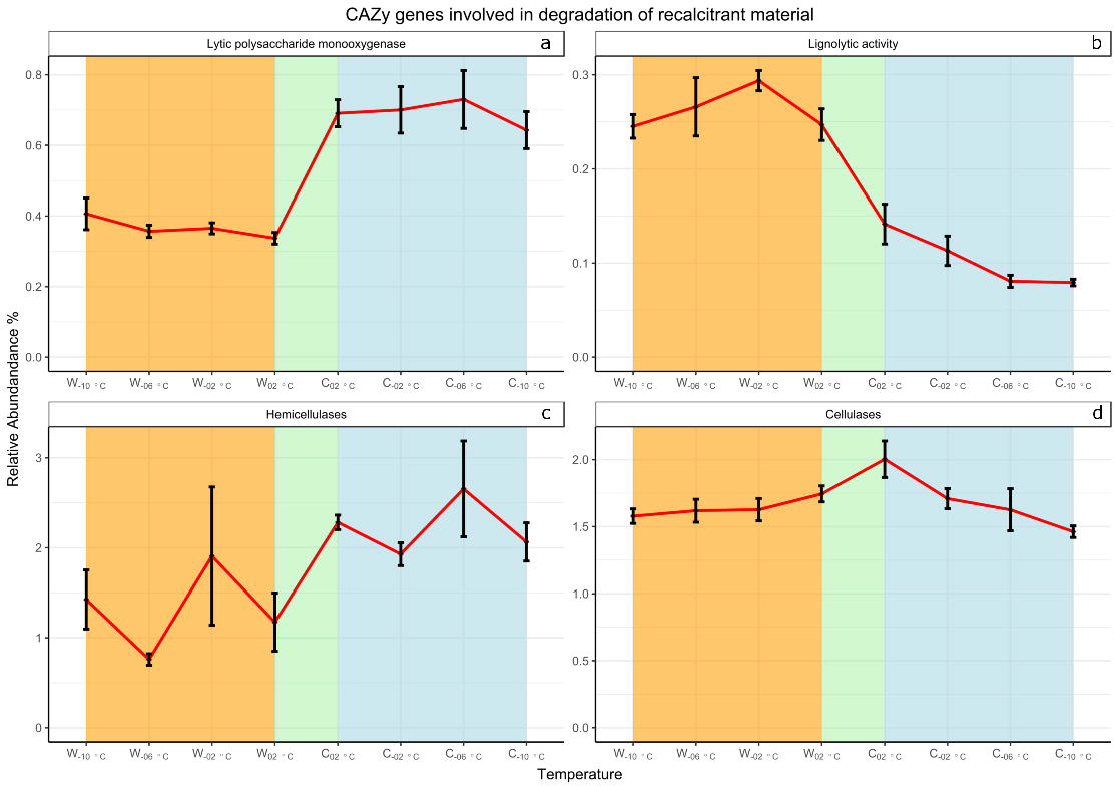
The relative abundance of transcripts annotated in CAZy as involved in degradation of complex plant and fungal material by encoding, a) lytic polysaccharide monooxygenases, b) lignolytic activity, c) hemicellulases, and d) cellulases. Error bars indicate standard error of the mean.

### Annotation of transcripts related to nitrogen cycling - alignment against NCycDB database

We detected a total of 233 annotated functions when aligning to the NCycDB database (a complete list of all gene functions can be found in Supporting Information Data Sheet S5). First, we investigated the response of the main soil nitrogen cycling pathways and, second, we investigated key gene families involved in soil nitrogen cycling. The major response of pathways related to nitrogen cycling occurred during the 16 days at 2 °C, when the relative abundance of transcripts related to ‘Organic degradation and synthesis’ and anaerobic ammonium oxidation (anammox) increased (*P* < 0.05), and transcripts related to assimilatory nitrate reduction, denitrification, and nitrification decreased (*P* < 0.05) (Fig. 5). We found no significant changes to transcripts assigned to nitrogen fixation. Several key nitrogen cycling gene families showed a change (*P* < 0.05) in the relative abundance of transcripts during the 16 days at 2 °C (Fig. 6). Thus, we observed a decrease in archaeal (but not bacterial) *amoA*, and denitrifier *narG* and *nirK*. In addition, we observed a response at the onset of freezing (between C_2°C_ and C-_2°C_) of several denitrifier transcripts (representing *narG*, *nirK*, *nirS* and *norB/C*). The NCycDB also includes genes encoding particulate methane monooxygenase, *pmoA*, *pmoB* and *pmoC*. Transcripts assigned to these genes decreased (*P* < 0.05) during the 16 days at 2 °C (Fig. 6C).

**Fig. 5.**
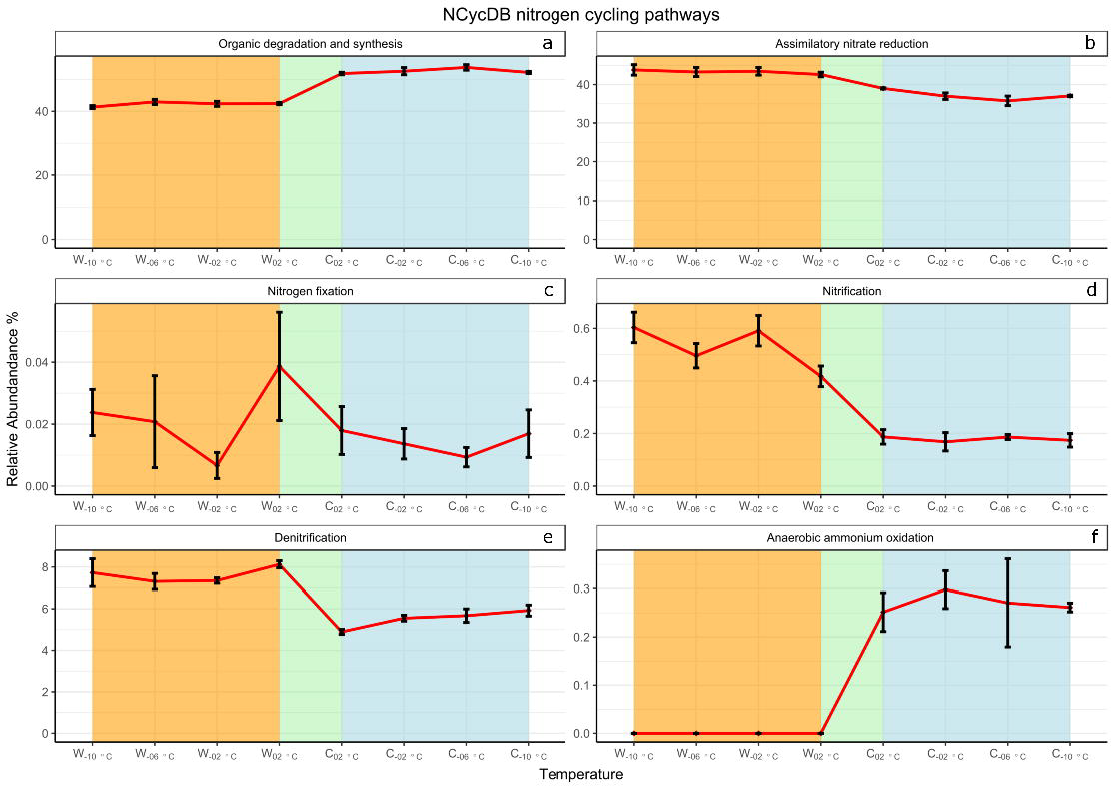
The relative abundance of transcripts annotated in NCycDB as involved in nitrogen cycling pathways, a) organic degradation and synthesis, b) assimilatory nitrate reduction, c) nitrogen fixation, d) nitrification, e) denitrification, and f) anaerobic ammonium oxidation. Error bars indicate standard error of the mean.

**Fig. 6.**
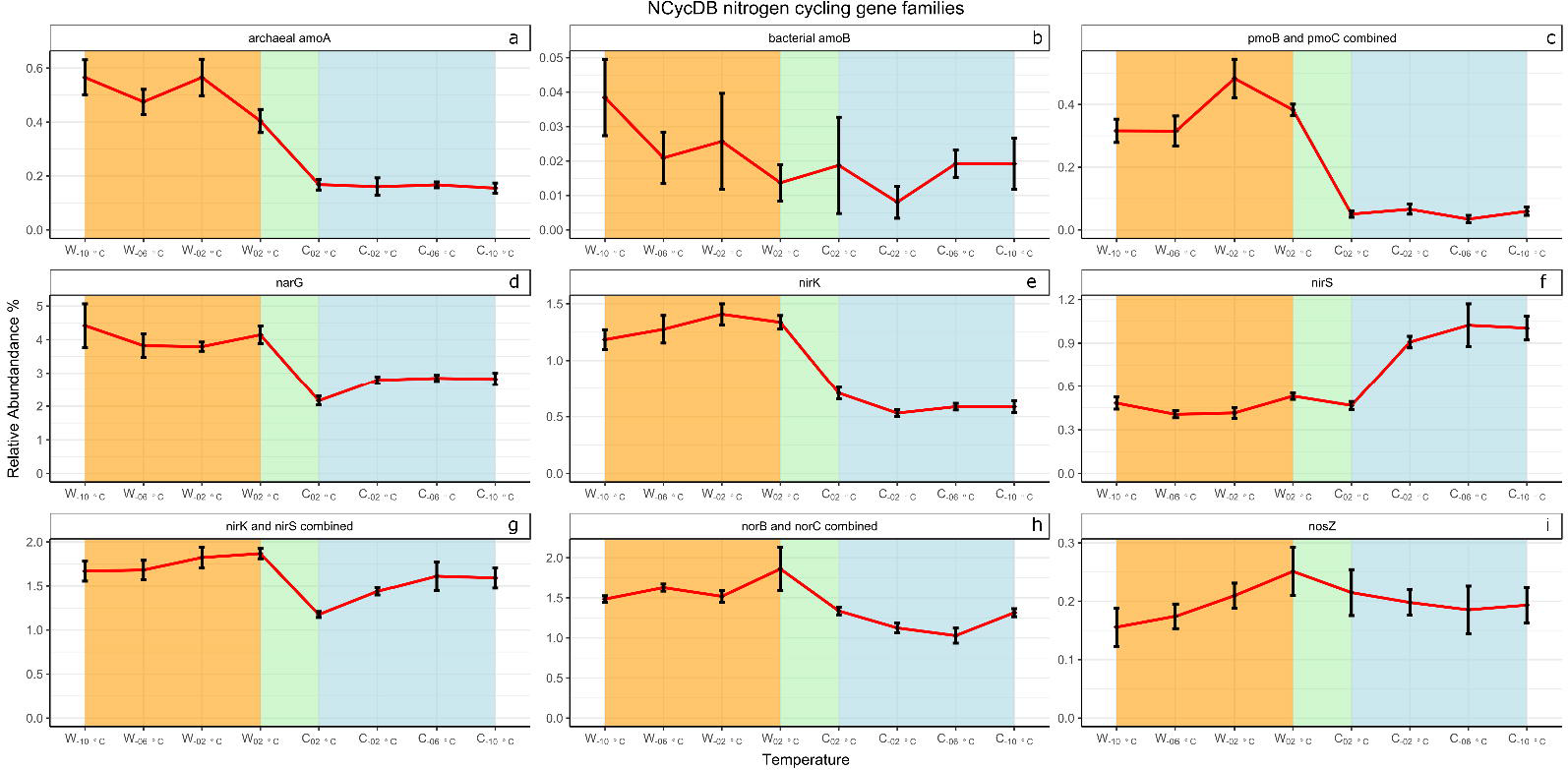
The relative abundance of transcripts annotated in NCycDB as involved in key nitrogen cycling and methane oxidation gene families, a) archaeal *amoA*, b) bacterial *amoA*, c) *pmoB* and *pmoC* combined, d) *narG*, e) *nirK*, f) *nirS*, g) *nirK* and *nirS* combined, h) *norB* and *norC* combined, and i) *nosZ*. Error bars indicate standard error of the mean.

## Discussion

To the best of our knowledge, this study was the first to investigate microbial gene expression during warming and subsequent cooling of an Arctic soil at several sub-zero temperatures and following freezing. Within both the warming and the cooling phase, gene expression was hardly affected by temperature change under frozen conditions. Also, we observed only moderate changes in gene transcription following transition between frozen (−2 °C) and thawed (+2 °C) states. However, following 16 days at 2 °C, we observed major transcriptional changes of genes involved in protein production, lignocellulose degradation, and nitrogen cycling.

### Transcription related to carbon cycling

We observed no changes in transcription of genes related to carbon metabolism during warming from −10 °C to 2 °C. However, between W_2°C_ and C_2°C_, we found an increase in transcription of genes in the eggNOG categories ‘Carbohydrate transport and metabolism’ and ‘Energy production and conversion’ as well as in CAZy functions related to degradation of lignocellulose and chitin. In accordance with the increase in the categories ‘Carbohydrate transport and metabolism’ and ‘Cell motility’ [reflecting enhanced bacterial and archaeal ability to access carbon and nutrient sources in the unfrozen soil], the concentration of dissolved organic carbon (DOC) decreased during the 16 days at 2 °C. The DOC pool mainly represents easily degradable and accessible organic matter. Thus, the DOC made available to microorganisms upon soil thaw may have contributed to initiate microbial activity and may have ‘kick started’ the breakdown of more complex/recalcitrant soil organic carbon (Coolen et al., 2011).

Complex soil organic carbon mainly consists of residues originating from plants or fungi (Clemmensen et al., 2013) and changes to the degradation of these residues in active layer soils are of great concern as temperatures in the Arctic are increasing (Schuur et al., 2008). We did not detect any transcripts related to degradation of plant and fungal polymers after going through all the annotated functions in the eggNOG dataset. Likewise, Coolen and Orsi (2015) did not detect hemicellulase, cellulase or laccase related transcripts before and after thawing of permafrost soil, while in an active layer peat soil from Svalbard, Tveit et al. (2014) assigned 0.02 – 0.08 % of transcripts to hemicellulase and cellulase genes. Depending on the database and workflow used we were only able to assign functions to 2.6 % of the filtered putative mRNA reads using the M5nr database and eggNOG annotation, while Tveit et al. (2014) were able to annotate 8 – 16 % of the putative mRNA reads matching the RefSeq database, and similar studies of a temperate forest soil assigned 28 % of all predicted coding regions to functional categories (Žifčáková et al. 2016) and of a temperate soil contaminated with copper assigned less than 9 % of the reads (Jacquiod et al., 2019). Our low annotation rates were likely partly due to limitations of the database when working with (Arctic) soils and partly to our stringent bioinformatic pipeline filtering less abundant contigs and potential non-coding RNAs. However, the pipeline adds more confidence to the annotation output (Anwar et al., 2019).

In contrast, annotation using CAZy revealed many transcripts involved in lignocellulose degradation. This was probably related to database specialization as CAZy is a well-curated and small database (database size influences the expected number of chance high-scoring segments and, hence, e-values), which uses more sensitive hidden Markov modelling for annotation. The increase in transcripts assigned by CAZy to encode lytic polysaccharide monooxygenase, cellulose, hemicellulose and chitinase genes at C_2°C_ indicates that soil thaw enhances degradation of soil organic polymers of plant or fungal origin. A pronounced response to thawing was associated with lytic polysaccharide monooxygenases, which are produced by numerous fungi and bacteria (Johansen, 2016). They constitute the first wave of attack on the most recalcitrant natural polysaccharides and initiate degradation of lignocellulose (Johansen, 2016) and chitin (Vaaje-Kolstad et al., 2010). Thus, lytic polysaccharide monooxygenases are secreted relatively early in the degradation process, while cellulases only appear at elevated levels later in the degradation process (Navarro et al., 2014).

The initiation of chitin, cellulose and hemicellulose gene transcription happened within 17 days of soil thawing and the resulting enzymes may have contributed to the observed increase in CO_2_ production. The soil microorganisms degrading lignocellulose and chitin seem to be able to respond to thaw within days and this response rate has implications for our understanding of CO_2_ emission from tundra soils. The microbial degraders of complex soil organic matter may be able to make the most of short summers and of freeze-thaw cycles that result in unfrozen conditions for days or weeks during autumn, winter and spring. In contrast, transcripts related to lignolytic gene activity decreased markedly upon soil thaw. This was likely caused by a decrease in fungal activity as the number of fungal 18S rRNA gene transcripts dropped significantly following thaw in the same experimental setup (Schostag et al., 2019). Fungi play a substantial role in the degradation of lignin in soils (Boer et al., 2005), including Arctic soils (Rinnan and Baath, 2009), but our experimental setup involving soil homogenization may have disrupted the activity of hyphae-forming fungi with lignin-degrading capabilities.

Transcripts of genes encoding particulate methane monooxygenase responsible for oxidation of methane at low concentration also decreased markedly from W_2°C_ to C_2°C_, which coincided with a shift in methane emission as the soil changed from a (small) net sink to a (small) net source of methane. The bacterial taxa performing methane oxidation in soils are generally slow growing (Islam et al., 2015) and we hypothesize that they were not able to outgrow the intense grazing by fast-growing protozoa initiated by soil thaw (Schostag et al., 2019) [the increase in the eggNOG category ‘Chromatin structure and dynamics’ likely reflects an increase in protozoan activity] leading to a decrease in numbers of methanotrophic bacteria. This may influence the ability of methanotrophic bacteria following spring thaw to oxidize atmospheric methane and methane emitted from deeper soil strata when these thaw.

### Transcription related to nitrogen cycling

Arctic plants and soil microorganisms are often limited by nitrogen availability (Elser et al., 2007) and the Arctic receives low rates of atmospheric nitrogen deposition (Dentener et al., 2006). This makes fixation of atmospheric nitrogen by cyanobacteria, either free-living or associated with mosses or lichenized fungi, the most important source of nitrogen to terrestrial Arctic ecosystems (Reed et al., 2011). We observed no transcriptional changes and only rather few transcripts related to nitrogen fixation. This may partly be explained by our experimental procedure, which excluded soil surface- and moss-associated cyanobacteria (mosses and the upper 1 cm of soil were removed) and included incubation in darkness. In addition, no leguminous plants are found at the site where the soil was sampled. This leaves few possibilities for nitrogen fixation in our experimental steup, being either *via* cyanobacteria or rhizobia.

Similar to transcripts related to carbon cycling, nitrogen cycling transcripts revealed the most pronounced changes following 17 days of thaw. The increase in transcripts involved in nitrogen-related ‘organic degradation and synthesis’ indicates i) enhanced degradation of nitrogen-containing organic matter as it coincided with the decrease in concentration of dissolved organic nitrogen (Tab. 1) and the increase in carbon cycling (Figs. 3A, 4A, C, D), and/or ii) enhanced synthesis of proteins and other nitrogen-containing compounds as the increase also coincided with that of protein production (Figs. 3C, D).

The pools of dissolved organic nitrogen and inorganic nitrogen showed pronounced changes during the experiment. The concentration of dissolved organic nitrogen decreased after 16 days at 2 °C likely due to microbial uptake and degradation leading to ammonification. Ammonification in combination with decreased nitrification led to an increase in soil ammonium concentration. Because transcription of genes assigned to nitrification decreased, the increase in nitrate concentration likely link to the decrease in transcripts assigned to nitrate-consuming processes, i.e. assimilatory nitrate reduction and denitrification. In Arctic soils, the former has been suggested to be more important for microbial nitrate consumption compared to denitrification (Taş et al., 2018), which was corroborated by the much larger number of transcripts assigned to assimilatory nitrate reduction compared to denitrification in our soil.

Archaeal *amoA* transcripts outnumbered bacterial *amoA* more than ten-fold when the soil thawed, and this dominance of archaeal ammonia-oxidizing transcriptional activity over bacterial is in accordance with other studies (Leininger et al., 2006; Alves et al., 2013; Feld et al., 2015). While the number of bacterial *amoA* transcripts was stable during the experiment, the number of archaeal *amoA* declined markedly following 16 days at 2 °C. The relative increase of bacterial nitrifiers over archaeal nitrifiers during the short Arctic summer before refreezing is likely due to the presence of released nutrients from dead organisms. In Feld et al. (2015), we previously found that archaeal *amoA* expression benefits less from release of nutrients in a soil that was fumigated compared to bacterial *amoA* expression. Nitrifying archaea are generally slow growing organisms (Könneke et al., 2005; Tourna et al., 2011) and in line with the discussion above on methane-oxidizing bacteria, we hypothesize that the nitrifying archaea were not able to outgrow the intense grazing by protozoa. Ammonium concentration increased substantially during the experiment and nitrifying bacteria are adapted to higher ammonium concentration compared to nitrifying archaea (Martens-Habbena et al., 2009). However, ammonium concentration in our samples was low in comparison to other Arctic soils (e.g. Alves et al. [2013] and Osborne et al. [2016]) and apparently not sufficiently high to enhance bacterial nitrifying activity following soil thaw. The decline in the activity of ammonia-oxidizing archaea following thaw may lower the potential nitrogen loss mediated by denitrification and leaching of nitrate and, hence, enhance availability of inorganic nitrogen to plants at the onset of the plant-growing season.

In contrast to nitrification, the number of transcripts assigned to anaerobic ammonium oxidation (anammox) increased following 16 days at 2 °C. This increase in anammox transcripts is dramatic considering the slow growth rate of these organisms (Kartal et al., 2013) and may be caused by increased ammonium and nitrite concentrations (the latter was not measured during the experiment). Little is known about anammox in Arctic soils, and the large increase in transcript numbers indicate that this process may be a hitherto overlooked source of nitrogen loss from Arctic soils.

Taş et al. (2018) found a high abundance of nitrifying organisms and a low genomic potential for denitrification across Arctic polygonal tundra soils. In contrast, we found a more than ten-fold higher abundance of transcripts assigned to denitrification compared to nitrification indicating a higher relative abundance of denitrification in our soil. Despite a build-up of soil nitrate between W_2°C_ and C_2°C_, the number of denitrification-associated transcripts decreased. This may be caused by the decrease in DOC concentration leading to enhanced competition from other heterotrophic bacteria and/or by changes in soil oxygen status (which we did not measure). In contrast to the other nitrogen and carbon cycle pathways, the number of transcripts assigned to the different steps in the denitrification pathway changed upon re-freezing (between C_2°C_ and C_−2°C_). Thus, the number of transcripts encoding nitrate and nitrite reductases increased, while a decrease was observed for nitric oxide reductase transcripts. Two different nitrite reductases are known, NirK encoded by *nirK* and NirS encoded by *nirS*. Individual denitrifying bacteria and archaea are only known to produce one of these reductases, as no denitrifying organism has been found that produces both. Strikingly, the number of transcripts assigned to *nirK* decreased upon re-freezing, while the number for *nirS* increased indicating an undescribed difference in the transcription of the genes encoding the two nitrite reductases e.g. in response to low temperature (Holtan-Hartwig et al., 2002), low water availability, and/or changes in oxygen status (Bakken et al., 2012).

The number of transcripts assigned to *nosZ* encoding the catalytic unit of nitrous oxide reductase was markedly lower than the number of transcripts assigned to the genes encoding the three other steps in the denitrification pathway. However, it is not possible to translate transcript numbers to enzyme activity, e.g. posttranscriptional assembly of *nosZ* is inhibited by low pH (Liu et al., 2014), and the low number of *nosZ* transcripts did not lead to high N_2_O emission rates. Throughout the experiment, the soil showed negligible net emission rates of N_2_O, which is common for Arctic soils (Christensen, 1999) probably because they are poised for nitrogen assimilation and not to release gaseous nitrogen compounds (Taş et al., 2018) reflecting a nitrogen-poor environment.

### Post-thawing stress responses

Only transcription of genes assigned to thirteen annotated eggNOG, one CAZy and four NCycDB functions were significantly up or downregulated one day after thawing. One of the initial responses to thaw was a possible stress response involving an increase in transcripts related to putative molecular chaperones, i.e. chaperonin GroEL (HSP60 family), that assist correct folding of proteins. In addition, the eggNOG category ‘Defence mechanisms’ decreased between W_2°C_ and C_2°C_. These represent a modest response to soil thaw compared to other studies (Mackelprang et al., 2011; Coolen & Orsi, 2015) and probably reflects the shorter time span between thawing and sampling in our experiment and our more robust annotation protocol (Anwar et al., 2019). The modest response to thaw indicates a lag phase longer than one day, which we initially hypothesized to be caused by the microorganisms focussing on downregulating non-stress genes as part of their stress response (Horn et al., 2007). However, we found little evidence that the stress response related to chaperone production caused a downregulation of non-stress related genes, as only five eggNOG and a single NCycDB function were downregulated one day after thaw.

The modest stress response may enable the microorganisms to focus on non-stress functions such as obtaining energy and nutrients from degradation of soil lignocellulose and chitin. Thus, the enhanced production of chaperones preceded a substantial increase in transcription of numerous genes following an additional 16 days at 2 °C, when 30 % of the annotated eggNOG functions were either significantly up or downregulated. A large fraction of the upregulated functions were related to transcription and translation, which was also observed in permafrost soil thawed for 11 days at 4 °C (Coolen & Orsi, 2015). Even though ribosomal numbers cannot be directly linked to an increase in microbial activity (Blazewicz et al., 2013), the fourfold increase in CO_2_ production rates during the 16 days at 2 °C indicates enhanced microbial activity. We think that the enhanced emission of CO_2_ was not due to release of previously trapped gas in ice, as the emission rate was measured 17–18 days after thaw and other experiments in our lab indicate that trapped CO_2_ is emitted from frozen soil within a few days of soil thaw (unpublished data).

The increase in transcripts related to translational activity also included molecular chaperones and ‘Cold shock proteins’. Chaperones have been found at high relative abundance in soil metaproteomic studies (Williams et al., 2010; Zampieri et al., 2016) and up to 60 % of identified proteins in an Alaskan active layer soil matched chaperones (Hultman et al., 2015). In our study, ‘Cold shock proteins’ were not induced upon or during freezing as has been observed before (Piette et al., 2011), but only following the 17-day period of stable temperature above freezing. Thus, the ‘Cold shock proteins’ were not part of a cold shock response *sensu stricto*, but were likely assisting protein folding as new proteins were produced or the changes in environmental conditions activated previously produced proteins. It has been suggested that changes in chaperone activity might be a direct microbial response to environmental fluctuations that have an effect on protein stability (Feder & Hofmann, 1999).

### Stress responses to re-freezing

Compared to permafrost soil, active layer soil contains a lower abundance of cold-shock proteins and other stress response genes (Yergeau et al., 2010; Mackelprang et al., 2011; Hultman et al., 2015). However, we hypothesized that re-freezing of the soil would elicit a cold-shock response as psychrophilic and cold-adapted microorganisms produce a large number of cold-shock proteins (De Maayer et al., 2014). The microorganisms in our soil responded to freezing by upregulating genes associated with 31 different eggNOG functions and downregulating genes associated with eight functions. Most of these were annotated as unknown functions, but we identified a decrease in transcription of genes related to cell motility upon freezing likely as a response to ice physically inhibiting prokaryotic motility. Also, two of the upregulated functions were associated with defense against oxidative stress, i.e. production of catalase [converting H_2_O_2_ to H_2_O and O_2_] and a DNA-binding ferritin-like protein denoted as an oxidative damage protectant. Elevated transcription of genes involved in production of catalases was seen in psychrophilic bacteria (Raymond-Bouchard et al., 2018) and genes involved in DNA repair was transcribed at higher abundance in frozen compared to thawed permafrost soil (Coolen & Orsi, 2015). Aerobic and facultative anaerobic microorganisms, including psychrophilic bacteria isolated from active layer soils (D’Amico et al., 2006; Mykytczuk et al., 2013), produce a number of enzymes that can neutralize reactive oxygen species (ROS) (Brioukhanov & Netrusov, 2007), which can damage DNA, proteins and cell membranes (Cabiscol et al., 2010; Ezraty et al., 2017). Oxidative stress may be enhanced at colder temperatures due to increased oxygen solubility (Weiss 1970; Sotelo et al., 1989) and higher enzyme activity (and hence production of ROS) initiated to adapt to reduced catalytic rates (Chattopadhyay et al., 2011; De Maayer et al., 2014). In contrast, diffusion rates of oxygen are lowered with decreasing temperature and ice formation effectively blocks transport of oxygen.

### Transcriptional response to warming or cooling at sub-zero temperatures

The minor transcriptional changes at sub-zero temperatures suggest that the microbial communities only responded marginally to either an increase or a decrease in temperature between −10 °C and −2 °C. The negligible response was not due to inactive microorganisms as CO_2_ production occurred at −6 °C and microbial communities in Arctic soils were shown to sustain activity and growth at similar or even lower temperatures (Panikov et al., 2006; Tuorto et al., 2014). We hypothesized that the temperature change spanning 8 °C would affect microbial transcriptional activity related to carbon metabolism, nitrogen cycling and stress, but at sub-zero temperatures, water availability and not temperature *per se* has the largest effect on microbial activity (Öquist et al., 2009; Tilston et al., 2010). The unfrozen water content in soils is influenced by salt concentration and organic matter composition (Drotz et al., 2010) while the unfrozen water content has been reported to show a small (Aanderud et al., 2013) or moderate (Panikov et al., 2006; Tilston et al., 2010) response to temperature changes occurring at the sub-zero temperatures employed in our experiment. We did not estimate water availability in the frozen soil samples, but the changes in water availability (and hence nutrient availability) between −10 °C and −2 °C may have been too small to elicit a detectable transcriptional response in our experimental setup.

Individual bacterial mRNAs have reported lifetimes from seconds to more than an hour (Condon, 2003; Deutscher, 2006), but in soil, *invA* mRNA was detected after 48 hours at 5 °C while it survived less than 4 hours at 15 °C or 25 °C (Garcia et al., 2010). These experiments were carried out at 5 °C or above and we have not been able to find information on the turnover of soil microbial mRNA at sub-zero temperatures. However, the long lifetime of *invA* in soil at 5 °C compared to at 15 °C and 25 °C (Garcia et al., 2010) indicates that mRNA may have extended lifetime at sub-zero temperatures. Thus, the minor transcriptional changes observed between −10 °C and −2 °C during warming and cooling may partly result from a slow turnover of mRNA at these temperatures.

## Conclusions

We detected only minor changes in the transcribed genes at sub-zero temperatures. Stress related transcripts, mainly defense against oxidative stress, were enhanced upon re-freezing of the soil, while no stress response was observed upon thawing. The transcriptional response to thawing was moderate after one day, but increased markedly after 17 days with a concomitant fourfold increase in CO_2_ production. This response was dominated by an increase in transcription of genes related to protein production and genes implicated in the degradation of soil cellulose, hemicellulose and chitin. In contrast, nitrogen cycle pathways were downregulated, except for anaerobic ammonium oxidation and degradation and synthesis of organic nitrogen. These findings may have implications for our understanding and modelling of carbon dioxide emission, nitrogen cycling and plant nutrient availability in Arctic soils.

## Supporting information

Supporting Information Data Sheet S1

Supporting Information Data Sheet S2

Supporting Information Data Sheet S3

Supporting Information Data Sheet S4

Supporting Information Data Sheet S5

## Methods

### Soil sample collection, preparation and incubation

An active layer soil core (0 – 14 cm soil depth; 8 cm diameter) was collected April 2014 in Sassendalen, Svalbard, Norway (latitude 78.270961, longitude 17.228315), using a core catcher with a motorized hand drill. The sampling site is characterized as a dry tundra dominated by *Dryas octopetala* and *Cassiope tetragona* (Vanderpuye et al., 2002; Elvebakk, 2005) with a mean annual temperature of −6 °C and a mean annual precipitation of ~200 mm water equivalents (Ingólfsson, 2011). The core was transported to our laboratory in Copenhagen, Denmark, in a styrofoam box with cooling elements at −18 °C. A temperature logger and visual inspection of the core indicated that it did not thaw during transport. In Copenhagen, all sample preparation was carried out in a −14 °C freeze lab before incubation. To avoid microbial contamination, all surfaces were sterilized by UV exposure and all equipment were washed with ethanol. The core was cut in half and the inner core soil was aseptically sampled with a sterilized bore head (16 mm diameter) from the newly exposed surfaces and homogenized by smashing the soil with a hammer while in a sterile plastic bag obtaining a grain size of 1 – 10 mm. For detailed description of the sample preparation, see Bang-Andreasen et al. (2017). Forty replicate 2-g subsamples were placed in 5-mL Eppendorf tubes and incubated in a custom-made temperature chamber based on a recirculating cooling liquid, for details see Schostag et al. (2019).

The samples were incubated for a total period of 26 days initiated with a pre-incubation step at −10 °C for 40 hours, then gradually increasing the temperature to 2 °C over five days, keeping a stable temperature of 2 °C for an additional 16 days, and finally a cooling phase with decreasing temperatures to −10 °C over five days. In this way, our experiment simulated a short Arctic spring, summer and autumn where temperature changes happen over time. Soil temperature measurements from a site in Adventdalen (ca. 10 km away) at a similar altitude showed that temperature at 1 cm soil depth is below −10 °C during winter and increases from - 10 to −2 °C within an average of 14 days during spring, which is somewhat slower than the five days employed in our experimental setup. Samples were obtained every 40 hours during the warming phase corresponding to −10, −6, −2 and 2 °C; henceforth denoted W_−10°C_, W_−6°C_, W_−2°C_, and W_2°C_, respectively (Fig. 1). No sampling was performed for 16 days until the cooling phase, during which samples were collected every 40 hours corresponding to 2, −2, −6 and −10 °C; denoted C_−10°C_, C_−6°C_, C_−2°C_, and C_2°C_, respectively. At each sampling point, five replicate subsamples were collected and immediately snap frozen in liquid nitrogen and stored at −80 °C until RNA isolation.

### Soil analyses

Soil physiochemical parameters were analyzed in a parallel set of 2-g soil samples incubated in 5-mL Eppendorf tubes as described above. These samples were incubated in separate tubes enabling us to snap freeze the subsamples for RNA isolation without additional handling of the soil. Due to limitations on the amount of soil that we could obtain from the soil core using our soil subsampling procedure we were not able to analyze soil physiochemical parameters and gas production rates at all eight time points. We selected four time points for the analysis of the physiochemical parameters: W_−6°C_, W_2°C_, C_2°C_, and C_−6°C_. At each time point, a total of 16 g soil was collected. For analysis of water extractable nutrients, 4 g soil was shaken (120 rpm) in 20 mL ddH_2_O for 10 min at 5 °C. All extracts were filtered through Whatman GF-D filters (Sigma-Aldridge, Copenhagen, Denmark) and frozen at −18 °C until analysis. A subset of the filtered extracts was used for pH measurements with a pH meter. Dissolved organic carbon (DOC) was analyzed with a Shimadzu TOC-L CSH/CSN total organic carbon analyzer (Shimadzu, Kyoto, Japan), while dissolved organic nitrogen (DON) was analyzed using a FIAstar 5000 (FOSS Tecator, Höganäs, Sweden) after digesting the extracts in 2 M HCl with selenium as a catalyst. Ammonium (NH_4_^+^) was analyzed using the indophenol blue method and nitrate (NO_3_^−^) colorimetrically using the cadmium reduction method, both with a FIAstar 5000 flow injection analyser (Foss, Hillerød, Denmark).

Carbon dioxide (CO_2_), methane (CH_4_) and nitrous oxide (N_2_O) production rates were measured by incubating 5 g soil in 50-mL serum flasks sealed with a butyl rubber stopper to which we added 15 mL of ambient air (to compensate for headspace being extracted from the flasks during sampling). To determine the gas production rates, 3-mL headspace samples were extracted at W_−6°C_, W_2°C_, and C_2°C_, and again 24 hours later. Gas samples were transferred to 3-mL Exetainer vials (LABCO, Lampeter, UK) and analyzed using an autosampler (Mikrolab Aarhus, Højbjerg, Denmark) and a 7890A GC system (Agilent Technologies, Glostrup, Denmark) equipped with a flame ionization detector and an electron capture detector.

### RNA isolation and cDNA sequencing

RNA isolation was carried out with 2 g of soil using RNA PowerSoil Total RNA Isolation Kit (Mo Bio Laboratories, Carlsbad, CA. USA) with phenol chloroform isoamyl alcohol 25:24:1 (Sigma-Aldrich) following the instructions of the manufacturer and resulting in 100 μL of RNA solution. DNase treatment on a 10-µL subsample was performed using DNase max (Mo Bio Laboratories) following the manufacturer’s instructions. The quality of DNase treated RNA was analyzed with Bioanalyser 2100 (Agilent Technologies, Glostrup, Denmark) and the quantity was estimated using Qubit 2.0 (Thermo Fisher Scientific, Life Technologies, Nærum, Denmark) with Qubit RNA HS Assay Kit. One hundred ng RNA from each sample were prepared for cDNA sequencing using NebNext Ultra Directional RNA Library Prep Kit for Illumina (New England Biolabs, Ipswich, MA, USA) according to the manufacturer’s instructions. Samples were sequenced using Illumina Hiseq 2500, rapid mode 150 bp paired-end, at the National High-throughput DNA Sequencing Centre, University of Copenhagen. Thus, our protocol did not involve mRNA enrichment using rRNA subtraction methods that may bias the relative abundance of different mRNA transcripts (Tveit *et al.*, 2014), and the protocol did not involve amplification of the cDNA.

### Bioinformatics and statistical methods

Cutadapt v. 1.9.1 (Martin, 2011) was used to trim adapters, poly-A tails, and filter reads shorter than 60 nucleotides or with phred score below 20. Putative mRNA reads were separated from the total pool of RNA by aligning and filtering all reads against Silva SSUref119 (removing 16S and 18S rRNA reads) and Silva LSURef119 (removing 23S and 28S rRNA reads) reference databases (Quast et al., 2013) using SortMeRNA v.2.1 (Kopylova et al., 2012) tool. Potential mRNA reads from all samples were pooled and assembled using trinity v.2.0.6 (Grabherr et al., 2011). From the resulting assembled contigs, non-coding RNA contigs were filtered by aligning contigs to the Rfam database v12.0 (Nawrocki et al., 2015) using cmsearch v1.1.1 (significance threshold e-value < 10^−3^). 72 % of the potential mRNA reads were assembled in contigs. BWA (Li et al., 2009) aligner was used to map back non-ribosomal RNA input sequences used for assembly to coding mRNA contigs. Due to variable sample resolution we normalized contigs by filtering the ones with relative expression lower than 1 % of the number of reads in the sample with least number of sequences. The open reading frames (ORFs) of the contigs were predicted using Transeq from EMBOSS (Rice et al., 2000) and were aligned using SWORD (Vaser et al., 2016) against the M5nr protein database (Wilke et al., 2012). The output was then parsed with in-house scripts written in Python where we selected the contigs with a significant e-value as base threshold. From the selected pool, the best hit for each contig was selected based on a combination of e-value and alignment length. For general gene annotation, M5nr aligned contigs were annotated against eggNOG hierarchical database v 4.5 (Jensen et al., 2008) using in-house Python scripts. For specific annotation of genes related to carbohydrate degradation, predicted ORFs were aligned against CAZy (Cantarel et al., 2009) database using SWORD and annotated against CAZy hierarchical annotation. Similarly, for annotation of genes related to nitrogen cycling, predicted ORFs were aligned and annotated against NCycDB (Tu et al., 2019); a manually created database using 100 % sequence identity. These comparisons resulted in three separate abundance tables, a generalist one with eggNOG orthologs and subsequent number of reads from each sample, a carbohydrate-degradation specific one with CAZy enzymes, and a nitrogen cycling specific one with NCycDB enzymes.

The beta diversity and multivariate analyses were done with the raw and non-rarefied contingency tables using the R software version 3.0.2 (R Development Core Team 2011) with the vegan (Oksanen et al., 2017) and ade4 (Dray & Dufour, 2007) packages. Principal Component Analysis (PCA) was performed after centre-scaling normalization. A pattern search was applied to the original PCAs by grouping replicates together in order to perform a between-group analysis (BGA). The statistical significance of the selected grouping factor was tested with a Monte-Carlo simulation involving 10,000 permutations. Complementing PERMANOVA tests were performed on the Euclidean distance profiles using 10,000 permutations in order to assess the significance of the tested factors.

To identify which eggNOG, CAZy and NCycDB genes, gene families or functional subsystems were significantly differentially expressed at different incubation times we used DESeq2 (Love et al., 2014) module of SarTools pipeline (Varet et al., 2016). This was done using parametric mean-variance and independent filtering of false discoveries with Benjamini-Hochberg procedure (*P* > 0.05) to adjust for type 1 error.

## Acknowledgements

The authors thank Pia Bach Jacobsen for help and technical support in the laboratory, Martin Asser Hansen for bioinformatics support, and Per Ambus and Vagn Roland Moser for soil analyses.

## Availability of data

Raw sequence data were deposited in the NCBI Sequence Read Archive and are accessible through accession number SRP124869.

## Authors’ contributions

M.D.S., C.S.J. and A.P. designed the study, S.F. collected the soil, M.D.S. conducted the experiment, M.Z.A. and M.D.S. performed bioinformatics and statistical analyses with help from S.J., L.M., C.L. and T.M.V., M.D.S., C.S.J., M.Z.A. and A.P. interpreted the results, and A.P. and M.D.S. wrote the manuscript with critical feedback from all of the coauthors.

## Ethics approval and consent to participate

Not applicable for this study

## Consent for publication

Not applicable for this study

## Supporting Information Data

**Supporting Information Figure S1** The relative abundance of transcripts assigned by CAZy database to glycoside hydrolase (GH) families GH18 and GH19 and presumed to be involved in degradation of chitin. Error bars indicate standard error of the mean.

**Supporting Information Table S1** Soil physiochemical characteristics measured prior to incubation. Data are average ± standard error of the mean, n = 5.

**Supporting Information Table S2** Sequence stats during bioinformatic processing.

**Supporting Information Data Sheet S1** List of all eggNOG functions that were significantly up or downregulated when comparing all sub-zero warming samples with W_2°C_ and number of reads assigned to these functions in the individual samples.

**Supporting Information Data Sheet S2** List of all eggNOG functions that were significantly up or downregulated when comparing W_2°C_ with C_2°C_ and number of reads assigned to these functions in the individual samples.

**Supporting Information Data Sheet S3** List of all eggNOG functions that were significantly up or downregulated when comparing C_2°C_ with all sub-zero cooling samples and number of reads assigned to these functions in the individual samples.

**Supporting Information Data Sheet S4** List of all CAZy gene functions and number of reads assigned to these functions in the individual samples.

**Supporting Information Data Sheet S5** List of all NCycDB gene functions and number of reads assigned to these functions in the individual samples.

