## Supplementary figures and images for "Transcriptomic responses to warming and cooling of an Arctic tundra soil microbiome"

### Supporting Information Figure S1

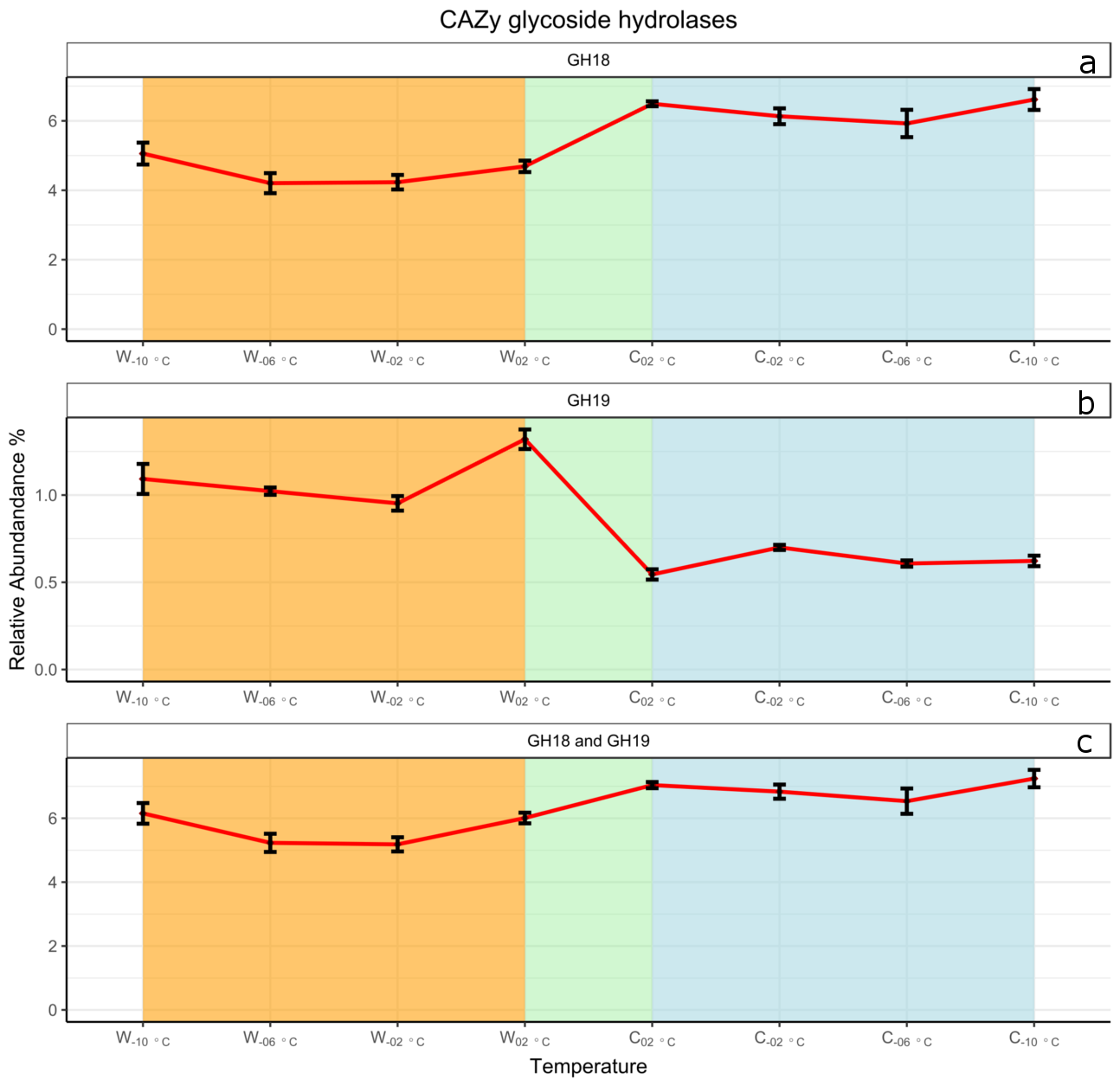
