## Supporting Information Table S1 for "Transcriptomic responses to warming and cooling of an Arctic tundra soil microbiome"

Supporting Information Table S1: Soil physiochemical characteristics measured prior to incubation. Data are average ± standard error of the mean, n = 5.

|  | % of dry soil |
| --- | --- |
| Water content | 39.6±1.4 |
| Total C | 7.19±0.32 |
| Total N | 0.48±0.02 |
| Clay (<2µm) | 3.1±0.4 |
| Silt (2-63µm) | 55.4±5.7 |
| Sand (>63µm) | 41.5±6.2 |
