## Supporting Information Table S2 for "Transcriptomic responses to warming and cooling of an Arctic tundra soil microbiome"

**Supporting Information Table S2 (part 1/2):** Sequence stats during bioinformatic processing.

|  |  |  | 1. Quality filtering and sorting | | | | | |  | 2. mRNA processing | | | | |
| --- | --- | --- | --- | --- | --- | --- | --- | --- | --- | --- | --- | --- | --- | --- |
| Samples | | | 1. HiSeq output |  | 1.2. Cutadapt |  | 1.3. SortMeRNA | |  | 2.1 Trinity Assembly | | |  | 2.2 BWA mapping^2^ |
| Sample | Temp°C | Event | # Raw sequences |  | # Sequences after QC |  | # SSU rRNA sequences | # Unaligned (mRNA) sequences |  | | Assembly stats^1^ (complete contig pool) |  |  | # Sequences mapped to contigs |
| Sample 9 | -10 | Thawing | 46183700 |  | 45948998 |  | 17382534 | 2270264 |  | | # contigs | 1,048,707 |  | 1646113 |
| Sample 12 | -10 | Thawing | 55492114 |  | 55383426 |  | 21430968 | 2824380 |  | | # contigs (>= 0 bp) | 1,048,707 |  | 2137885 |
| Sample 18 | -10 | Thawing | 34470870 |  | 34402890 |  | 13037252 | 1891948 |  | | # contigs (>= 1000 bp) | 8778 |  | 1411531 |
| Sample 39 | -10 | Thawing | 30665984 |  | 30550358 |  | 11577372 | 1694758 |  | | # contigs (>= 5000 bp) | 35 |  | 1216293 |
| Sample 2 | -6 | Thawing | 47320360 |  | 46866966 |  | 18099234 | 2475368 |  | | # contigs (>= 10000 bp) | 2 |  | 1923596 |
| Sample 4 | -6 | Thawing | 51021612 |  | 50777416 |  | 19676336 | 2384104 |  | | Largest contig | 10,795 |  | 1782598 |
| Sample 10 | -6 | Thawing | 27412914 |  | 27275256 |  | 10257690 | 1450622 |  | | Total length | 325,508,965 |  | 1067694 |
| Sample 23 | -6 | Thawing | 26248830 |  | 25921712 |  | 10150214 | 1364170 |  | | Total length (>= 0 bp) | 325,508,965 |  | 999339 |
| Sample 3 | -2 | Thawing | 40234324 |  | 39940428 |  | 15386908 | 1980858 |  | | Total length (>= 1000 bp) | 325,508,965 |  | 1467220 |
| Sample 5 | -2 | Thawing | 33713442 |  | 33456648 |  | 12749510 | 1768074 |  | | Total length (>= 5000 bp) | 228487 |  | 1297735 |
| Sample 20 | -2 | Thawing | 31906624 |  | 31521648 |  | 12162300 | 2039410 |  | | Total length (>= 10000 bp) | 20807 |  | 1449284 |
| Sample 32 | -2 | Thawing | 25618252 |  | 25526726 |  | 9784920 | 1455372 |  | | N50 | 695 |  | 1052700 |
| Sample 1 | 2 | Thawing | 30836820 |  | 30238088 |  | 11273534 | 1688770 |  | | N75 | 573 |  | 1201505 |
| Sample 7 | 2 | Thawing | 39027706 |  | 38740938 |  | 14817250 | 1830098 |  | | L50 | 26213 |  | 1326614 |
| Sample 16 | 2 | Thawing | 26763020 |  | 26676470 |  | 10091252 | 1355542 |  | | L75 | 47499 |  | 956506 |
| Sample 40 | 2 | Thawing | 24878052 |  | 24807942 |  | 9425930 | 1298108 |  | | GC (%) | 56.97 |  | 895542 |
| Sample 6 | 2 | Freeze | 45859720 |  | 45582796 |  | 17989118 | 1640840 |  | |  |  |  | 1150127 |
| Sample 26 | 2 | Freeze | 31947380 |  | 31683100 |  | 12154320 | 1494600 |  | |  |  |  | 978273 |
| Sample 30 | 2 | Freeze | 34006324 |  | 33675514 |  | 13396254 | 1397478 |  | |  |  |  | 980565 |
| Sample 36 | 2 | Freeze | 29919812 |  | 29780700 |  | 11861974 | 1152818 |  | |  |  |  | 798302 |
| Sample 17 | -2 | Freeze | 30280296 |  | 30215094 |  | 11589856 | 1328908 |  | |  |  |  | 937239 |
| Sample 27 | -2 | Freeze | 45663492 |  | 44972226 |  | 17883170 | 1946880 |  | |  |  |  | 1370395 |
| Sample 37 | -2 | Freeze | 27470464 |  | 27367006 |  | 11129916 | 1057484 |  | |  |  |  | 729485 |
| Sample 41 | -2 | Freeze | 27688494 |  | 27627060 |  | 11054822 | 991724 |  | |  |  |  | 724048 |
| Sample 11 | -6 | Freeze | 36180912 |  | 36050476 |  | 13813996 | 1424434 |  | |  |  |  | 995824 |
| Sample 21 | -6 | Freeze | 32637904 |  | 32376936 |  | 12732506 | 1326702 |  | |  |  |  | 957815 |
| Sample 31 | -6 | Freeze | 28202622 |  | 27790404 |  | 11061472 | 1188602 |  | |  |  |  | 868889 |
| Sample 38 | -6 | Freeze | 28505990 |  | 28414418 |  | 10923686 | 1218392 |  | |  |  |  | 817047 |
| Sample 15 | -10 | Freeze | 46404696 |  | 46285026 |  | 17462248 | 1973596 |  | |  |  |  | 1404250 |
| Sample 19 | -10 | Freeze | 46289146 |  | 46209680 |  | 17399460 | 1865614 |  | |  |  |  | 1260929 |
| Sample 22 | -10 | Freeze | 30228794 |  | 29946866 |  | 11952712 | 1277730 |  | |  |  |  | 933587 |
| Sample 34 | -10 | Freeze | 43611752 |  | 43320124 |  | 17052614 | 1789518 |  | |  |  |  | 1322835 |
|  |  | **Total:** | **1,136,692,422** |  | **1,129,333,336** |  | **436,761,328** | **52,847,166** |  |  | |  |  | **38,061,765** |

**Supporting Information Table S2 (part 2/2):** Sequence stats during bioinformatic processing.

| Samples | | | 3.3. Normalization | 3.4. Rfam filtering | 3.5. SWORD | 3.6. SWORD | 3.7. SWORD |
| --- | --- | --- | --- | --- | --- | --- | --- |
| Sample | Temp°C | Event | # Sequences remaining | # Sequences remaining | # Sequences annotated to eggNOG functional subsystems | # Sequences annotated to CAZy enzymes | # Sequences annotated to NCycDB enzymes |
| Sample 9 | -10 | Thawing | 1189272 | 851029 | 20575 | 51508 | 13947 |
| Sample 12 | -10 | Thawing | 1554047 | 1179678 | 29320 | 65840 | 16032 |
| Sample 18 | -10 | Thawing | 1051066 | 774286 | 19783 | 40536 | 11296 |
| Sample 39 | -10 | Thawing | 898706 | 647713 | 15515 | 36523 | 9584 |
| Sample 2 | -6 | Thawing | 1502382 | 1117319 | 24528 | 58017 | 14820 |
| Sample 4 | -6 | Thawing | 1298263 | 964082 | 28749 | 53489 | 13906 |
| Sample 10 | -6 | Thawing | 797885 | 572635 | 16385 | 31440 | 8199 |
| Sample 23 | -6 | Thawing | 737650 | 521758 | 14351 | 32383 | 8609 |
| Sample 3 | -2 | Thawing | 1069007 | 779934 | 21676 | 43875 | 11692 |
| Sample 5 | -2 | Thawing | 949062 | 698444 | 18730 | 40262 | 11500 |
| Sample 20 | -2 | Thawing | 1095828 | 774897 | 17067 | 40728 | 10857 |
| Sample 32 | -2 | Thawing | 794708 | 561900 | 13467 | 29481 | 7891 |
| Sample 1 | 2 | Thawing | 868196 | 613006 | 16071 | 35861 | 9965 |
| Sample 7 | 2 | Thawing | 947324 | 701384 | 16051 | 43908 | 12785 |
| Sample 16 | 2 | Thawing | 678962 | 496070 | 14267 | 31785 | 8934 |
| Sample 40 | 2 | Thawing | 626392 | 449816 | 11491 | 28432 | 8934 |
| Sample 6 | 2 | Freeze | 813132 | 561519 | 11394 | 75614 | 16508 |
| Sample 26 | 2 | Freeze | 702634 | 458625 | 10865 | 54870 | 13257 |
| Sample 30 | 2 | Freeze | 728474 | 477974 | 11648 | 52038 | 12617 |
| Sample 36 | 2 | Freeze | 594633 | 383952 | 8953 | 44001 | 10142 |
| Sample 17 | -2 | Freeze | 688374 | 466932 | 11564 | 53237 | 13165 |
| Sample 27 | -2 | Freeze | 1010300 | 666354 | 16972 | 98883 | 25915 |
| Sample 37 | -2 | Freeze | 521658 | 351526 | 7701 | 45566 | 11759 |
| Sample 41 | -2 | Freeze | 533867 | 360829 | 8889 | 43100 | 11369 |
| Sample 11 | -6 | Freeze | 716455 | 482586 | 13386 | 70691 | 16272 |
| Sample 21 | -6 | Freeze | 698973 | 497194 | 12228 | 67236 | 17734 |
| Sample 31 | -6 | Freeze | 659953 | 424692 | 10290 | 54568 | 14662 |
| Sample 38 | -6 | Freeze | 583113 | 392369 | 10857 | 63599 | 14562 |
| Sample 15 | -10 | Freeze | 1023220 | 679218 | 22072 | 91552 | 22089 |
| Sample 19 | -10 | Freeze | 874156 | 623549 | 16794 | 85045 | 22394 |
| Sample 22 | -10 | Freeze | 706738 | 468509 | 13043 | 54813 | 13812 |
| Sample 34 | -10 | Freeze | 1002190 | 646901 | 16653 | 86700 | 21667 |
|  |  |  | 27,916,620 | 19,646,680 | 501,336 | 1,705,581 | 435,733 |
|  |  |  | 78,015 | 49,324 | Functional COGs  652 | CAZy Enzymes  1648 | NcycDB genes  233 |
